# Length–length and length–weight relationships for four small pelagic fishes in the Kuroshio–Oyashio current system

**DOI:** 10.1101/2020.12.16.423010

**Authors:** Sho Furuichi, Yasuhiro Kamimura, Ryuji Yukami

**Author notes:** **Correspondence**: Sho Furuichi, Fisheries Resources Institute, Japan Fisheries Research and Education Agency, Yokohama, Japan.

## Abstract

Length–length relationships and length–weight relationships were estimated for four small pelagic fishes (Japanese sardine *Sardinops melanostictus*, Japanese anchovy *Engraulis japonicus*, chub mackerel *Scomber japonicus*, and spotted mackerel *Scomber australasicus*) in the Kuroshio–Oyashio current system. Fish samples were collected from surface–midwater trawl surveys and commercial purse-seine fisheries between September and October of 2020 in the western North Pacific. Total length (TL), fork length (FL), and standard length (SL) were measured to 0.01 cm, and whole body weight (W) was measured to the nearest 0.01 g for each individual. All length–length relationships (TL–FL, TL–SL, and FL–SL) were highly significant (*p* < 0.0001), with *r*^2^ > 0.98 in all species. Length–weight relationships (W–TL, W–FL, and W–SL) were also highly significant (*p* < 0.0001), with *r*^2^> 0.98 in all species. This study provides a useful reference for biological studies and stock assessments of these small pelagic fishes.

## 1. INTRODUCTION

Length–length relationships (LLRs) and length–weight relationships (LWRs) of fishes are key tools in fish biology and ecology, fisheries science, and fisheries management (Froese, 2006; Froese et al., 2011). Length–length relationships are needed to convert length measurements from one length type into another and are also important in comparative growth studies. Length–weight relationships are used to predict weight from length and to calculate production and biomass of a fish population in stock assessment.

In the Kuroshio–Oyashio current system (around the Japanese Archipelago), the small pelagic fish community is dominated by Japanese sardine *Sardinops melanostictus* Temminck & Schlegel, 1844, Japanese anchovy *Engraulis japonicus* Temminck & Schlegel, 1846, chub mackerel *Scomber japonicus* Houttuyn, 1782, and spotted mackerel *Scomber australasicus* Cuvier, 1832. The ecological and commercial importance and remarkable fluctuations in the abundances of these small pelagic fishes have inspired studies ranging in scope from the dynamics of single stocks to the dynamics of small pelagic fish communities at ecosystem and global scales (Chavez, 2003; Oozeki et al., 2019). Although numerous researchers from different institutions and disciplines have studied the same ecosystem, and even the same fish stocks, these researchers have often used different standards for fish length measurement (i.e., standard length [SL], fork length [FL], or total length [TL]) (e.g., Jung et al., 2008; Kang et al., 2009; Yukami et al., 2008). Therefore, conversion equations (that is, LLRs) for these various length measurement standards are required for comparative or integrative research among different geographic regions and species. However, there is currently little information on LLRs for the four small pelagic fishes in the Kuroshio–Oyashio current system. In addition, although LWRs for these fishes have been reported, studies have been limited to specific geographic regions and length-measurement standards.

In this study, we provide LLRs and LWRs among TL, FL, and SL for the four small pelagic fish species (*S. melanostictus*, *E. japonicus*, *S. japonicus*, and *S. australasicus*) that dominate the Kuroshio–Oyashio current system.

## 2. MATERIALS AND METHODS

Specimens were obtained from a trawl survey and from commercial purse-seine catches. In the trawl survey, 35 surface–midwater trawls were conducted in the western North Pacific off northern Japan and the Kuril Islands (38°20′N‒46°58′N, 142°8′E‒174°50′E) during September–October 2020. A surface–midwater trawl net (opening dimensions: 30 m × 30 m; cod-end mesh size: 17.5 mm) was towed once at each station (15–60 min duration at 3.5–5.0 kn [1.8–2.6 m/s], depths < 30–40 m). At each station, fish species were identified on board, and up to 380 individuals of each species were randomly selected and stored at –25 °C. Samples from purse-seine catches were collected from commercial landings at the Hachinohe Port (an important fishing port in northern Japan) in October 2020. All sampled fish were measured to the nearest 0.01 cm in standard length (SL), fork length (FL), and total length (TL) in the laboratory. Total weight was measured to the nearest 0.01 g.

To estimate LLRs, linear relationships between the various measurement types (TL, FL, and SL in cm) were derived by using major-axis (MA) regression. MA regression is used to find the axis that minimizes errors in both measured variables, and is particularly useful for comparative and integrative study as it allows bi-directional conversions.

LWRs were estimated by using the equation: *W* = *a* × *L*^b^, where *W* is total body weight (g), *L* is length (TL, FL, or SL in cm), *a* is the coefficient of proportionality, and *b* is the coefficient of allometry. Parameters *a* and *b* were estimated by non-linear least-squares regression. Additionally, 95% confidence intervals of *a* and *b*, and the coefficient of determination (*r*^2^) were calculated. All statistical analyses were performed in R version 3.6.3 (R Development Core Team, 2020).

## 3. RESULTS

Sample sizes, body-length and body-weight ranges, and estimated LWR and LLR parameters for each fish species are presented in Tables 1, 2, and 3. All LLRs were highly significant (*p* < 0.0001), with *r*^2^ values >0.98. All LWRs were also highly significant (*p* < 0.0001), with *r*^2^ values >0.94.

**Table 1.** Sample sizes and ranges of body length and weight of four small pelagic fish species caught in the western North Pacific.

| Species | *n* | TL (cm) |  | FL (cm) |  | SL (cm) |  | W (g) |  |
| --- | --- | --- | --- | --- | --- | --- | --- | --- | --- |
|  |  | Min | Max | Min | Max | Min | Max | Min | Max |
| *Sardinops melanostictus*<br>(Temminck & Schlegel, 1844) | 1809 | 12.5 | 25.2 | 11.4 | 22.6 | 10.9 | 21.5 | 12.9 | 142.4 |
| *Engraulis japonicus*<br>(Temminck & Schlegel, 1846) | 664 | 4.7 | 16.2 | 4.3 | 15.4 | 4.0 | 14.4 | 0.43 | 28.6 |
| *Scomber japonicus*<br>(Houttuyn, 1782) | 3682 | 6.3 | 46.1 | 5.9 | 43.0 | 5.3 | 39.8 | 1.6 | 874.9 |
| *Scomber australasicus*<br>(Cuvier, 1832) | 312 | 15.7 | 34.4 | 14.6 | 32.2 | 13.6 | 29.4 | 31.2 | 369.5 |

Abbreviations: *n*, sample size; TL, total length; FL, fork length; SL, standard length; W, total body weight.

**Table 2.** Estimated length–length relationship parameters for four small pelagic fish species in the western North Pacific.

| Species | Equation | $a$ (95% CI) | $b$ (95% CI) | $r^2$ |
| --- | --- | --- | --- | --- |
| *Sardinops melanostictus*<br>(Temminck & Schlegel, 1844) | TL = $a + b$ (FL) | −0.241 (−0.327 to −0.156) | 1.09 (1.09 to 1.10) | 0.99 |
| | TL = $a + b$ (SL) | 0.371 (0.295 to 0.447) | 1.13 (1.12 to 1.13) | 0.99 |
| | FL = $a + b$ (SL) | 0.559 (0.485 to 0.633) | 1.03 (1.03 to 1.04) | 0.99 |
| *Engraulis japonicus*<br>(Temminck & Schlegel, 1846) | TL = $a + b$ (FL) | 0.121 (0.064 to 0.178) | 1.07 (1.06 to 1.08) | 0.99 |
| | TL = $a + b$ (SL) | 0.358 (0.307 to 0.409) | 1.12 (1.11 to 1.13) | 0.99 |
| | FL = $a + b$ (SL) | 0.221 (0.167 to 0.276) | 1.05 (1.04 to 1.05) | 0.99 |
| *Scomber japonicus*<br>(Houttuyn, 1782) | TL = $a + b$ (FL) | −0.136 (−0.177 to −0.095) | 1.07 (1.07 to 1.08) | 0.99 |
| | TL = $a + b$ (SL) | −0.050 (−0.094 to −0.007) | 1.16 (1.16 to 1.16) | 0.99 |
| | FL = $a + b$ (SL) | 0.081 (0.049 to 0.113) | 1.08 (1.08 to 1.08) | 0.99 |
| *Scomber australasicus*<br>(Cuvier, 1832) | TL = $a + b$ (FL) | −0.006 (−0.294 to 0.278) | 1.07 (1.05 to 1.08) | 0.98 |
| | TL = $a + b$ (SL) | 0.158 (−0.124 to 0.436) | 1.14 (1.13 to 1.16) | 0.98 |
| | FL = $a + b$ (SL) | 0.157 (−0.051 to 0.362) | 1.07 (1.06 to 1.08) | 0.99 |

Abbreviations: TL, total length; FL, fork length; SL, standard length; CI, confidence interval; $r^2$ , coefficient of determination.

**Table 3.** Estimated length–weight relationship parameters for four small pelagic fish species in the western North Pacific.

| Species | Equation | $a$ (95% CI) | $b$ (95% CI) | $r^2$ |
| --- | --- | --- | --- | --- |
| *Sardinops melanostictus*<br>(Temminck & Schlegel, 1844) | $W = a \times TL^b$ | 0.002 (0.002 to 0.003) | 3.45 (3.41 to 3.48) | 0.94 |
| | $W = a \times FL^b$ | 0.002 (0.002 to 0.003) | 3.54 (3.50 to 3.58) | 0.94 |
| | $W = a \times SL^b$ | 0.004 (0.004 to 0.005) | 3.4 (3.37 to 3.44) | 0.95 |
| *Engraulis japonicus*<br>(Temminck & Schlegel, 1846) | $W = a \times TL^b$ | 0.004 (0.003 to 0.004) | 3.2 (3.16 to 3.24) | 0.98 |
| | $W = a \times FL^b$ | 0.006 (0.006 to 0.007) | 3.08 (3.04 to 3.12) | 0.97 |
| | $W = a \times SL^b$ | 0.008 (0.007 to 0.008) | 3.10 (3.06 to 3.13) | 0.98 |
| *Scomber japonicus*<br>(Houttuyn, 1782) | $W = a \times TL^b$ | 0.007 (0.006 to 0.007) | 3.09 (3.08 to 3.10) | 0.99 |
| | $W = a \times FL^b$ | 0.008 (0.007 to 0.008) | 3.11 (3.10 to 3.12) | 0.99 |
| | $W = a \times SL^b$ | 0.010 (0.010 to 0.010) | 3.10 (3.09 to 3.11) | 0.99 |
| *Scomber australasicus*<br>(Cuvier, 1832) | $W = a \times TL^b$ | 0.006 (0.005 to 0.007) | 3.15 (3.09 to 3.21) | 0.96 |
| | $W = a \times FL^b$ | 0.007 (0.006 to 0.008) | 3.15 (3.09 to 3.20) | 0.97 |
| | $W = a \times SL^b$ | 0.006 (0.006 to 0.007) | 3.25 (3.20 to 3.39) | 0.98 |

Abbreviations: W, total body weight; TL, total length; FL, fork length; SL, standard length; CI, confidence interval; $r^2$ , coefficient of determination.

## 4. DISCUSSION

In the present study, we estimated LLRs for *S. melanostictus*, *E. japonicus*, *S. japonicus*, and *S. australasicus*. This is the first comprehensive report of LLRs for these four small pelagic fish species in the Kuroshio–Oyashio current system. Although these species have been studied in various parts of the world, previous studies employed a variety of standards for fish length measurements, making comparisons between and among studies difficult. Our results will be helpful for comparative or integrative research among different geographic regions and species.

In addition to LLRs, we also estimated LWRs for the four species. The allometric coefficients (*b*) of the LWRs were within the expected range of 2.5–3.5 (Froese, 2006), although there was some variability among species. This indicates that our estimates are plausible. However, we were not able to sample specimens from across a broad temporal and spatial range. LWRs can be affected by various factors such as season and location (Jellyman et al., 2013; Valle et al., 2020). Therefore, a broader sampling scheme in future studies could provide more robust LWRs.

In conclusion, our results provide new information on LLRs and LWRs for four small pelagic fish species in the Kuroshio–Oyashio current system. These data will be useful for further biological study and management of the four species.

## ACKNOWLEDGEMENTS

We are very grateful to Michiyo Ichikawa, Aki Narisawa, Akiko Sugawara, Yuko Saiki, and Tomiko Nagai, who greatly contributed to the data collection. We thank all those who helped in the collection of samples. We also thank English-speaking professional editors from ELSS, Inc., for English proofreading. Funding for this study was provided by the Japan Fisheries Research and Education Agency and the Fisheries Agency of Japan.

## CONFLICTS OF INTEREST

The authors declare no conflicts of interest.

## DATA AVALABILITY STATEMENT

The data used to support the findings of this study are available from the corresponding author upon request.

## Notes

### Competing Interest Statement

The authors have declared no competing interest.

